# Interpreting UniFrac with Absolute Abundance: A Conceptual and Practical Guide

**DOI:** 10.1101/2025.07.18.665540

**Authors:** Augustus Pendleton, Marian L. Schmidt

**Affiliations:** Department of Microbiology, Cornell University, 123 Wing Dr, Ithaca, NY 14850, USA

**Keywords:** Microbial Ecology, Beta Diversity, Absolute Abundance, Bioinformatics, UniFrac

## Abstract

*β*-diversity is central to microbial ecology, yet commonly used metrics overlook changes in microbial load, limiting their ability to detect ecologically meaningful shifts. Popular for incorporating phylogenetic relationships, UniFrac distances currently default to relative abundance and therefore omit important variation in microbial abundances. As quantifying absolute abundance becomes more accessible, integrating this information into *β*-diversity analyses is essential. Here, we introduce *Absolute UniFrac* (*U*^*A*^), a variant of Weighted UniFrac that incorporates absolute abundances. Using simulations and a reanalysis of four 16S rRNA metabarcoding datasets (from a nuclear reactor cooling tank, the mouse gut, a freshwater lake, and the peanut rhizospere), we demonstrate that Absolute UniFrac captures microbial load, composition, and phylogenetic relationships. While this can improve statistical power to detect ecological shifts, we also find Absolute Unifrac can be strongly correlated to differences in cell abundances alone. To balance these effects, we also incorporate absolute abundance into the generalized extension (*GU*^*A*^) that has a tunable *α* parameter to adjust the influence of abundance and composition. Finally, we benchmark *GU*^*A*^ and show that although computationally slower than conventional alternatives, *GU*^*A*^ is comparably insensitive to realistic noise in load estimates compared to conventional alternatives like Bray-Curtis dissimilarities, particularly at lower *α*. By coupling phylogeny, composition, and microbial load, Absolute Unifrac integrates three dimensions of ecological change, better equipping microbial ecologists to quantitatively compare microbial communities.

## Main Text

Microbial ecologists routinely compare communities using *β*-diversity metrics derived from relative abundances. Yet this approach overlooks a critical ecological dimension: microbial load. High-throughput sequencing produces compositional data, in which each taxon’s abundance is constrained by all others [1]. However, quantitative profiling studies show that cell abundance, not only composition, can drive major community differences [2]. In low-biomass samples, relying on relative abundance can allow contaminants to appear biologically meaningful despite absolute counts too low for concern [3].

Sequencing-based microbiome studies therefore rely on relative abundance even when the hypotheses of interest implicitly concern absolute changes in biomass. This creates a mismatch between ecological framing (growth, bloom magnitude, pathogen proliferation, disturbance recovery, or colonization pressure) and the information the *β*-diversity metric encodes [4]. As a result, *β*-diversity is often treated as if it includes biomass, even when absolute abundance is either not measured at all or is measured but excluded from the calculation (as in conventional UniFrac). Many studies therefore test biomass-linked hypotheses using a metric that normalizes biomass away [4]. A conceptual correction is needed in which *β*-diversity is understood as variation along three axes: composition, phylogeny and absolute abundance.

Absolute microbial load measurements are now increasingly obtainable through flow cytometry, qPCR, and genomic spike-ins, allowing biomass to be quantified alongside taxonomic composition [4, 5]. These approaches improve detection of functionally relevant taxa and mitigate the compositional constraints imposed by sequencing [1, 2]. Most studies that incorporate absolute abundance counts currently rely on Bray-Curtis dissimilarity, which can capture load but does not consider phylogenetic similarity [5–7]. Unifrac distances provide the opposite strength, incorporating phylogeny but discarding absolute abundance, as they are restricted to relative abundance data by construction [8]. This leaves no phylogenetically informed *β*-diversity metric that operates on absolute counts, despite the fact that biomass is central to many ecological hypotheses.

To evaluate the implications of incorporating absolute abundance into phylogenetic *β*-diversity, we use both simulations and reanalysis of four 16S rRNA amplicon datasets. The simulations use a simple four-taxon community with controlled abundance shifts to directly compare both Bray-Curtis and UniFrac in their relative and absolute forms, illustrating how each metric responds when abundance, composition, or evolutionary relatedness differ. We then reanalyze four real-world datasets spanning nuclear reactor cooling water [5], the mouse gut [3], a freshwater lake [9], and rhizosphere soil [10]. These differ widely in richness, biomass range, and ecological context, allowing us to test when absolute abundance changes align with or diverge from phylogenetic turnover. Together, these simulations and reanalyses provide the empirical foundation for interpreting Absolute UniFrac relative to existing *β*-diversity measures across the three axes of ecological difference: abundance, composition, and phylogeny.

We also extend Absolute UniFrac as was proposed by [11] to Generalized Absolute UniFrac that incorporates a tunable ecological dimension, *α*, and evaluate its impact across simulated and real-world datasets. As *α* increases, *β*-diversity is increasingly correlated with differences in absolute abundance, allowing researchers to fine tune the relative weight their analyses place on microbial load versus composition.

### Defining Absolute UniFrac

The UniFrac distance was first introduced by Lozupone & Knight (2005) and has since become enormously popular as a measure of *β*-diversity within the field of microbial ecology [12]. A benefit of the UniFrac distance is that it considers phylogenetic information when estimating the distance between two communities. After first generating a phylogenetic tree representing species (or amplicon sequence variants, “ASVs”) from all samples, the UniFrac distance computes the fraction of branch-lengths which is *shared* between communities, relative to the total branch length represented in the tree. UniFrac can be both unweighted, in which only the incidence of species is considered, or weighted, wherein a branch’s contribution is weighted by the proportional abundance of taxa on that branch [8]. The Weighted UniFrac is derived:

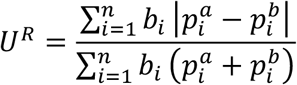

where the contribution of each branch length, *b*_*i*_, is weighted by the difference in the relative abundance of all species (*p*_*i*_) descended from that branch in sample *a* or sample *b*. Here, we denote this distance as *U*^*R*^, for “Relative UniFrac”. Popular packages which calculate weighted Unifrac—including the diversity-lib QIIME plug-in and the R packages phyloseq and GUniFrac—run this normalization by default [11, 13, 14].

Importantly, *U*^*R*^ is most sensitive to changes in abundant lineages, which can sometimes obscure compositional differences driven by rare to moderately-abundant taxa [11]. To address this weakness, Chen et al. (2012) introduced the generalized UniFrac distance (*GU*^*R*^), in which the impact of abundant lineages can be mitigated by decreasing the parameter *α*:

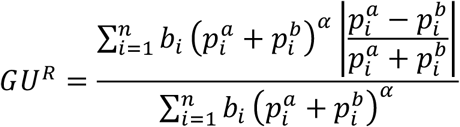

where *α* ranges from 0 (close to Unweighted UniFrac) up to 1 (identical to *U*^*R*^, above). However, if one wishes to use absolute abundances, both *U*^*R*^ and *GU*^*R*^ can be derived without normalizing to proportions:

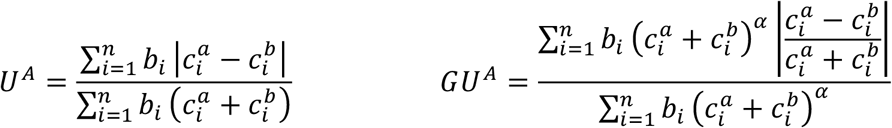

Where 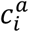 and 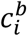 denote for the absolute counts of species descending from branch *b*_*i*_ in communities *a* and *b*, respectively. We refer to these distances as *Absolute UniFrac* (*U*^*A*^)and *Generalized Absolute Unifrac* (*GU*^*A*^). Although substituting absolute for relative abundances is mathematically straightforward, we found no prior work that examines UniFrac in the context of absolute abundance, either conceptually or in application. Incorporating absolute abundances introduces a third axis of ecological variation: beyond differences in composition and phylogenetic similarity, *U*^*A*^ also captures divergence in microbial load. This makes interpretation of *U*^*A*^ nontrivial, particularly in complex microbiomes.

### Demonstrating *β*-diversity metrics’ behavior with a simple simulation

To clarify how *U*^*A*^ behaves relative to existing *β*-diversity metrics, we first constructed a simple four-taxon simulated community arranged in a small phylogeny (Fig. 1A). By varying the absolute abundance of each ASV (1, 10, or 100), we generated 81 samples and 3,240 pairwise comparisons. For each pair, we computed four dissimilarity metrics: Bray-Curtis with relative abundance (*BC*^*R*^), Bray-Curtis with absolute abundance (*BC*^*A*^), Weighted UniFrac with relative abundance (*U*^*R*^), and Weighted UniFrac with absolute abundance (*U*^*A*^). These comparisons help to illustrate how each axis of ecological difference—abundance, composition, and phylogeny—is expressed by the different metrics.

**Figure 1.**
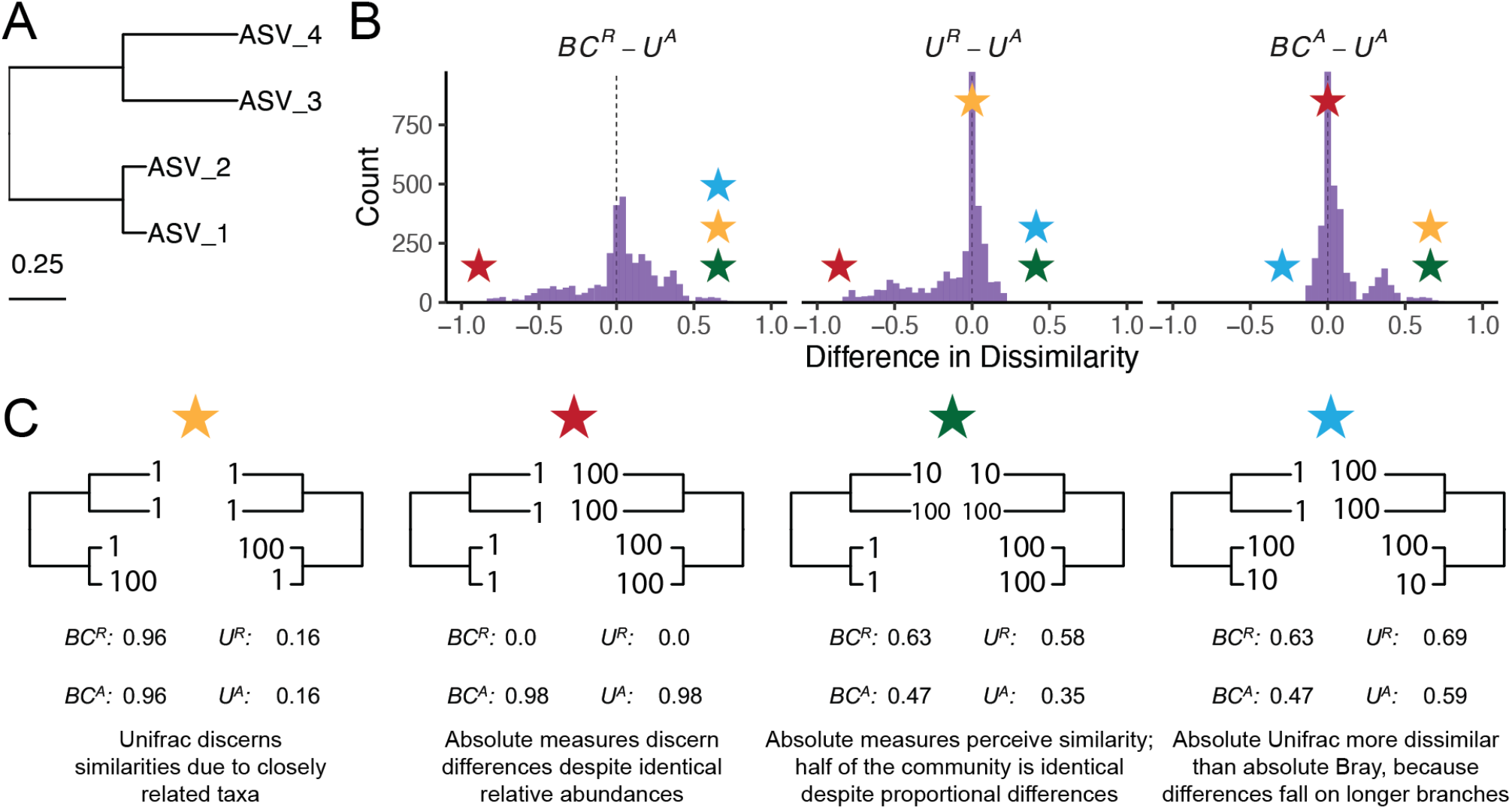
Simulated communities reveal how absolute abundance affects phylogenetic and non-phylogenetic β-diversity measures. (A) We constructed a simple four-ASV community with a known phylogeny and generated all permutations of each ASV having an absolute abundance of 1, 10, or 100, resulting in 81 unique communities and 3,240 pairwise comparisons. (B) Distributions of pairwise differences between weighted UniFrac using absolute abundance (*U*^*A*^) and three other metrics: Bray-Curtis using relative abundance (*BC*^*R*^), weighted UniFrac using relative abundance (*U*^*R*^), and Bray-Curtis using absolute abundance (*BC*^*A*^). (C) Illustrative sample pairs demonstrate how absolute abundances and phylogenetic structure interact to increase or decrease dissimilarity across metrics. Stars indicate where each scenario falls within the distributions shown in panel B. Actual values for each metric are displayed beneath each scenario.

*U*^*A*^ does not consistently yield higher or lower distances compared to other metrics, but instead varies depending on how abundance and phylogeny intersect (Fig. 1B). In the improbable scenario that all branch lengths are equal, *U*^*A*^ is always less than or equal to *BC*^*A*^ (Fig. S1). These comparisons emphasize that once branch lengths differ, incorporating phylogeny and absolute abundance alters the structure of the distance space. The direction and magnitude of that change depend on which branches carry the abundance shifts. *U*^*A*^ is also usually smaller than *BC*^*A*^ and is more strongly correlated with *BC*^*A*^ (Pearsons *r* = 0.82, *p* < 0.0001) than with *BC*^*R*^ (*r* = 0.41) and *U*^*R*^ (*r* = 0.55).

To better understand how these metrics diverge, we examined individual sample pairs (Fig. 1C). Scenario 1 (gold star) illustrates the classic advantage of UniFrac: ASV_1 and ASV_2 are phylogenetically close, so *U*^*R*^ and *U*^*A*^ discern greater similarity between samples than *BC*^*R*^ and *BC*^*A*^, which ignore phylogenetic structure. Scenario 2 (red star) highlights a limitation of relative metrics: two samples with identical relative composition but a 100-fold difference in biomass appear identical to *BC*^*R*^ and *U*^*R*^, but not to their absolute counterparts. In Scenario 3 (green star), incorporating absolute abundance decreases dissimilarity. *BC*^*A*^ and *U*^*A*^ are lower than their relative counterparts because half the community is identical in absolute abundance, even though their proportions differ. In contrast, Scenario 4 (blue star) shows that *U*^*A*^ can exceed *BC*^*A*^ when abundance differences occur on long branches, amplifying phylogenetic dissimilarity.

These scenarios demonstrate that *U*^*A*^ integrates variation along three ecologically relevant axes: composition, phylogenetic similarity, and microbial load, rather than isolating any single dimension. Because a given *U*^*A*^ value can reflect multiple drivers of community change, interpreting it requires downstream analyses to disentangle the relative contributions of these three axes. To evaluate how this plays out in real systems we next reanalyzed four previously published datasets spanning diverse microbial environments.

### Application of Absolute UniFrac to Four Real-World Microbiome Datasets

To illustrate the sensitivity of *U*^*A*^ to both variation in composition and absolute abundance, we re-analyzed four previously published datasets from diverse microbial systems. These datasets included: (i) nuclear reactor cooling water sampled across three reactor cycle phases [5]; (ii) the mouse gut sampled across multiple regions and between two diets [3]; (iii) depth-stratified freshwater communities sampled across two months [9]; and (iv) peanut rhizospheres under two crop rotation schemes (conventional rotation (CR) vs. sod-based rotation (SBR)), plant maturities, and irrigation treatments [10]. Absolute abundance was quantified by flow cytometry for the cooling water and freshwater datasets, droplet digital PCR for the mouse gut, and by qPCR for the rhizosphere dataset. Together these span a wide range of richness–from 215 ASVs in the cooling water to 24,000 ASVs in the soil–and microbial load–from as low as 4×10^5^ cells/ml (cooling water) up to 2×10^12^ 16S rRNA copies/gram (mouse gut). Additional details of the re-analysis workflow, including ASV generation and phylogenetic inference, are provided in the Supporting Methods.

We first calculated four β-diversity metrics for all sample pairs in each dataset and compared them to *U*^*A*^ (Fig. 2). The degree of concordance between *U*^*A*^ and other metrics was highly context dependent. In the cooling water dataset, *U*^*A*^ closely tracked all three alternatives, whereas in the remaining systems it diverged substantially. *U*^*A*^ also spanned a similar or wider range of distances than the other metrics. For example, in the soil dataset *BC*^*R*^ and *U*^*R*^ occupied a narrow range relative to the broad separation observed under *U*^*A*^.

**Figure 2.**
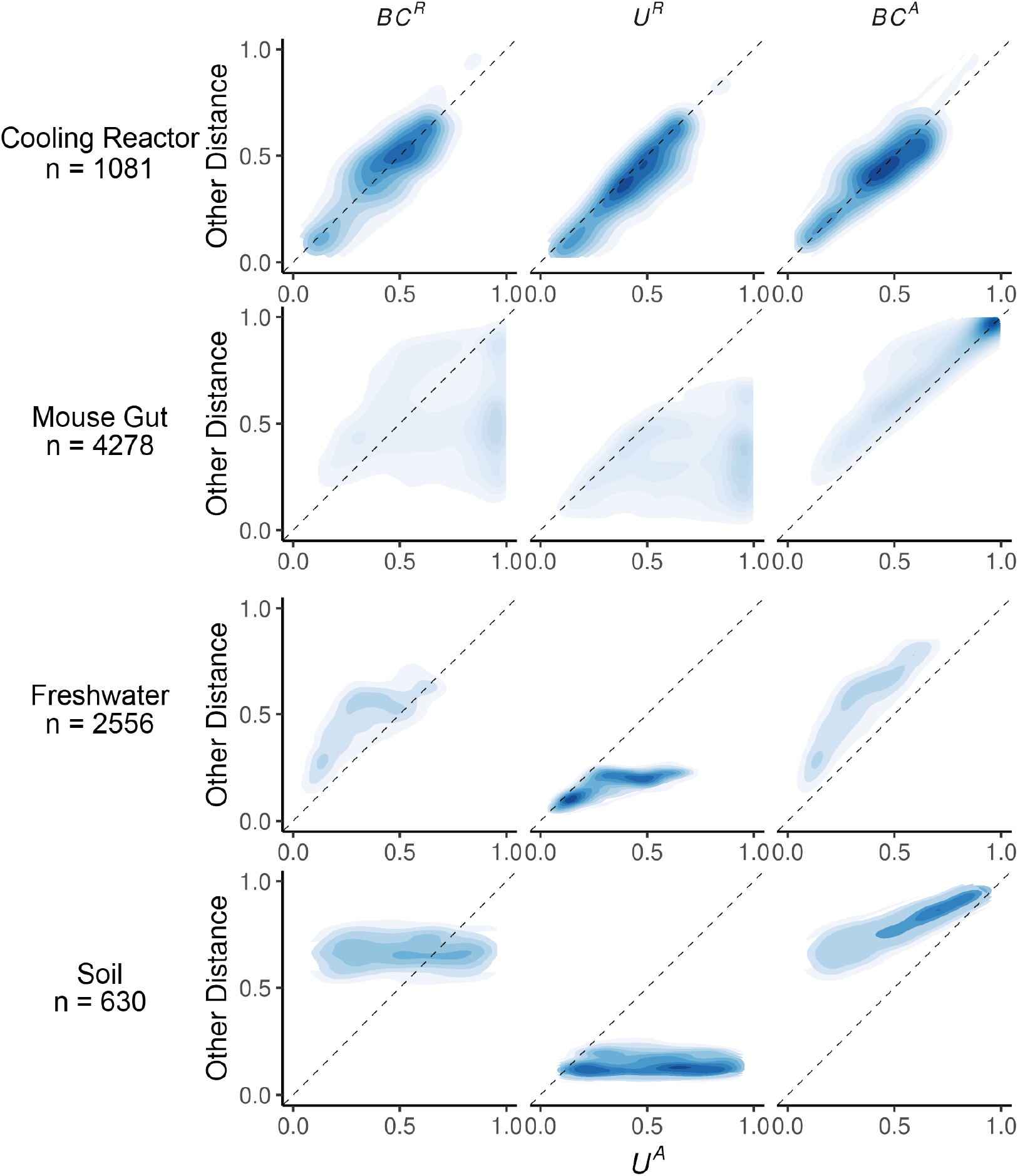
Absolute UniFrac (U^A^) compared with other β-diversity metrics across four real microbial datasets. Each panel shows pairwise sample distances for *U*^*A*^ (x-axis) against another metric (y-axis): Bray-Curtis using relative abundance (*BC*^*R*^, first column), weighted UniFrac using relative abundances (*U*^*R*^, second column), and Bray-curtis using absolute abundances (*BC*^*A*^, third column). Contours indicate the relative density of pairwise comparisons (n shown for each dataset, left), with darker shading corresponding to more observations. The dashed line marks the 1:1 relationship. Points above the line indicate cases where *U*^*A*^ is smaller than the comparator metric, while points below the line indicate cases where *U*^*A*^ is larger.

*U*^*A*^ generally reported distances that were similar to or greater than *U*^*R*^, consistent with the simulations shown in Fig. 1B. This reflects the ability of *U*^*A*^ to discern dissimilarity due to differences in microbial load, even when community composition is conserved. In contrast, *U*^*A*^ yielded distances that were similar to or lower than *BC*^*A*^, again matching the simulated behavior in Fig 1B. In these cases, phylogenetic proximity between abundant, closely-related ASVs leads *U*^*A*^ to register greater similarity than *BC*^*A*^.

Given these differences, we next quantified how well each metric discriminates among categorical sample groups. For each dataset, we ran PERMANOVAs to measure the proportion of variance (*R*^*2*^) and statistical power (*pseudo-F, p-*value) attributable to group structure, using groupings that were determined to be significant in the original publications. To evaluate how strongly absolute abundance contributed to this discrimination, we also calculated *GU*^*A*^ across a range of *α* values. The resulting *R*^*2*^ values are displayed in Fig. 3A, with the corresponding *pseudo-F* statistics and *p*-values provided in Fig. S2.

**Figure 3.**
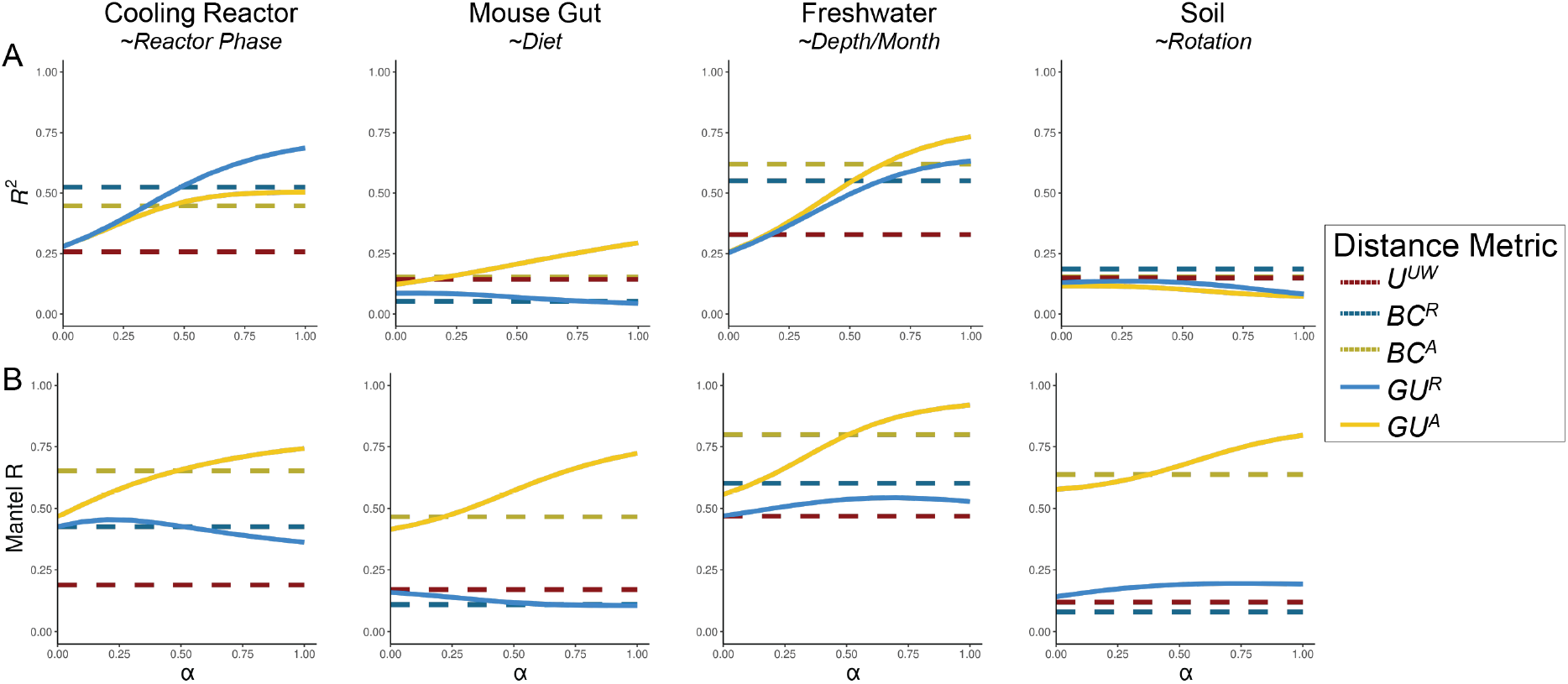
Discriminatory performance of U^A^ and related metrics across four microbial systems. (A) PERMANOVAs were used to quantify the percent variance (*R*^*2*^) explained by predefined categorical groups (shown in italics beneath each dataset name), with 1,000 permutations. PERMANOVA results were evaluated across five metrics and, where applicable, across eleven *α* values (0-1 in 0.1 increments). For consistency with the original studies, only samples from Reactor cycle 1 were used for the cooling-water dataset, only stool samples for the mouse gut dataset, and only mature rhizosphere samples for the dataset. (B) Mantel correlation (R) between each distance metric and the pairwise differences in absolute abundance (cell counts or 16S copy number), illustrating the degree to which each metric is driven by biomass differences.

As in Fig. 2, the performance of each metric was strongly context dependent (Fig. 3A). In the mouse gut and freshwater datasets, absolute-abundance-aware metrics (*BC*^*A*^ and *GU*^*A*^) explained the greatest proportion of variance (*R*^*2*^), and *R*^*2*^ generally increasing as *α* increased. In contrast, relative metrics captured more variation in the cooling water dataset (again at higher *α*), and all metrics explained comparably little variance in the soil dataset. Taken at face value, these trends might suggest that higher *α* values typically improve group differentiation.

However, this comes with a major caveat: at high *α* values, *GU*^*A*^ becomes strongly correlated with total cell count alone (Fig. 3B). Mantel tests confirmed that absolute-abundance metrics are far more sensitive to differences in microbial load than their relative counterparts. This behavior is intuitive, and to some extent desirable, because these metrics are designed to detect changes in microbial load even when composition remains constant. Yet at *α* = 1, *U*^*A*^ approaches a proxy for sample absolute abundance itself. In ordination space (Fig. S3), this can cause Axis 1 to correlate with absolute abundance and in some cases (freshwater and soil) produces strong horseshoe effects [15], potentially distorting ecological interpretation.

We recommend calibrating *α* based on research goals, modulating this effect by using *GU*^*A*^ across a range of *α* rather than relying on *U*^*A*^ (*α* = 1). Researchers should consider how much emphasis they want their dissimilarity metric to place on microbial load (Box 1). When biomass differences are central to the hypothesis being tested (for example, detecting cyanobacterial blooms), high *α* are recommended. In contrast, if microbial load is irrelevant or independent of the hypothesis in question, low *α* (or *U*^*R*^) may be preferred; for example in the soil dataset, fine-scale differences in composition may be obscured by random variation in microbial load.

In many systems, microbial biomass is one piece of the story, likely correlated to other variables being tested. If the importance of microbial load in the system is unknown, one potential approach is to calculate correlations as demonstrated in Fig. 3B and select an *α* prior to any ordinations or statistical testing. Again, correlation to cell count is valuable and is intrinsic to absolute abundance-aware measures, especially when microbial load is relevant to the hypotheses being tested. Correlations to cell count in *BC*^*A*^, an accepted approach in the literature, ranged from ∼0.5 up to ∼0.8. As a general recommendation from these analyses, we recommend *α* values in an intermediate range from 0.1 up to 0.6, wherein *GU*^*A*^ has similar correlation to cell count as *BC*^*A*^.

### Computational and Methodological Considerations

Applying *GU*^*A*^ in practice raises several considerations related to sequencing depth, richness, and computational cost. Both Bray-Curtis and UniFrac can be sensitive to sequencing depth because richness varies with read count [16–18]. Methods to address these concerns, including rarefaction, remain debated and are challenging to benchmark rigorously [19]. We do not evaluate the sensitivity of *U*^*A*^ or *GU*^*A*^ to different rarefaction strategies here, but we do provide a workflow and code for how we incorporated rarefaction into our own analyses (Box 2 and available code). This approach minimizes sequencing-depth biases while preserving abundance scaling for downstream *β*-diversity analysis.

*GU*^*A*^ is slower to compute than both *BC*^*A*^ and *U*^*R*^ because it must traverse the phylogenetic tree to calculate branch lengths for each iteration. Computational time of *GU*^*A*^ increases quadratically with ASV number and is 10-20 times slower, though up to 50 times slower, than *BC*^*A*^ (Fig. 4A-B). The number of samples or *α* values, however, have relatively little effect on runtime (Fig. S4). For a dataset of 24,000 ASVs, computing on a single CPU is still reasonable (1.4 minutes), but repeated rarefaction increases runtime substantially because branch lengths are redundantly calculated with each iteration. Allowing branch-length objects to be cached or incorporated directly into the GUnifrac workflow would considerably improve computational efficiency.

**Figure 4.**
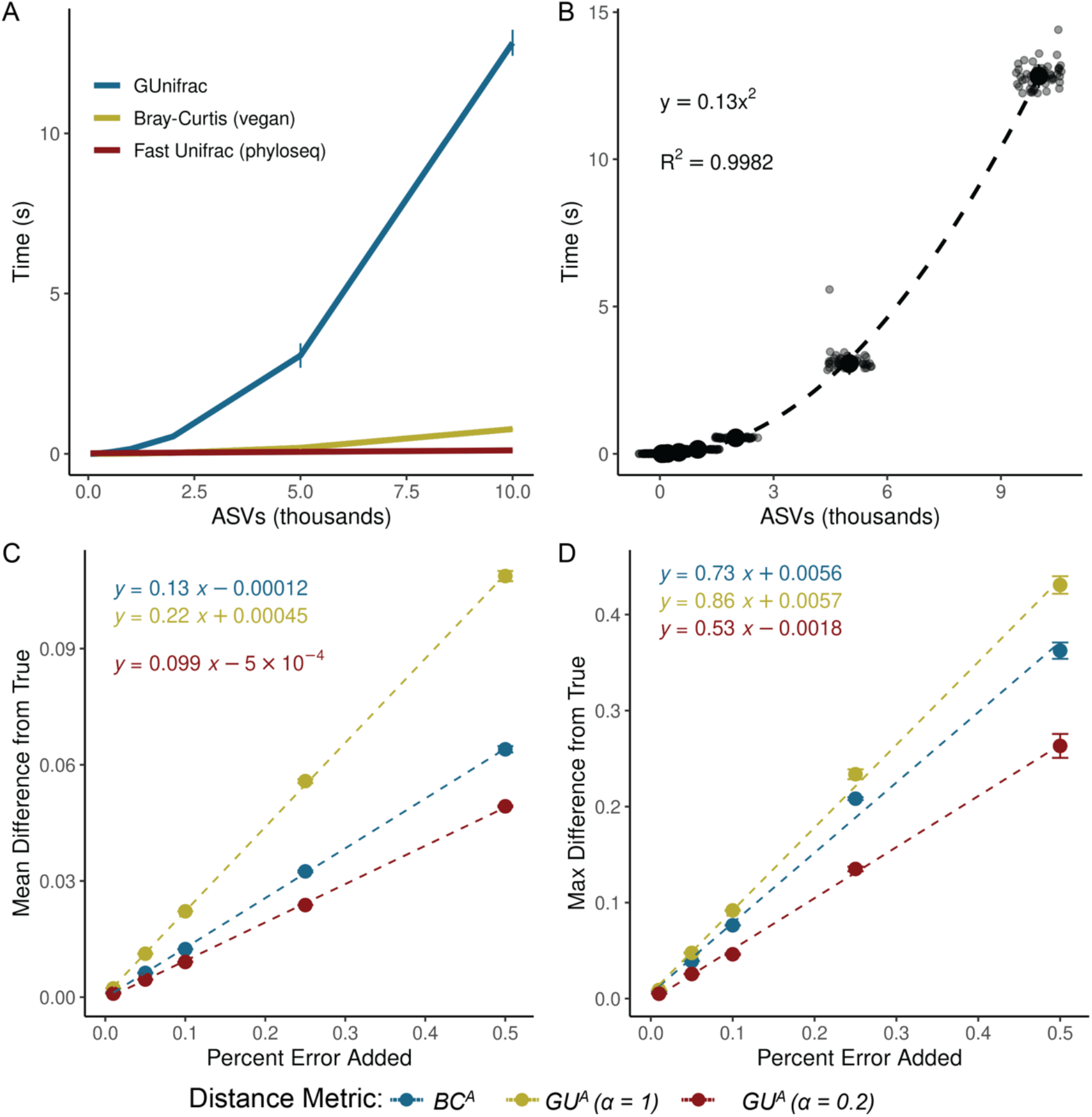
GU^A^ requires more computational time but remains resilient to quantification error. (A) Runtime for *GU*^*A*^ (GUniFrac package), *U*^*R*^ (FastUniFrac in the phyloseq package) and *BC*^*A*^ (vegan package) was benchmarked across 50 iterations on a sub-sampled soil dataset (mature samples only) [10], using increasing ASV richness (50, 100, 200, 500, 1,000, 2,000, 5,000, 10,000), with 10 samples and one *α* value per run (unweighted UniFrac is also calculated by default). Error bars represent standard deviation. (B) Quadratic relationship between *GU*^*A*^ computation time and ASV richness. Large center point represents median across 50 iterations, error bars (standard deviation) are too small to be seen. (C-D) Sensitivity of *GU*^*A*^ (*α* = 1 and 0.2) and *BC*^*A*^ to measurement error was evaluated by adding random variation (±1% to ±50%; Supporting Methods) to 16S copy number estimates in stool samples from the mouse gut dataset [3]. For each error level, 50 replicate matrices were generated and compared to the original values. Panels reflect the (C) mean difference and (D) max difference between the error-added metrics compared to the originals. Error bars represent the standard deviation of the average mean and max difference across 50 iterations.

We also evaluated the sensitivity of *GU*^*A*^ to measurement error in absolute abundance due to uncertainty arising from the quantification of cell number of 16S copy number. To assess the sensitivity of *GU*^*A*^ and *BC*^*A*^ to measurement error, we added random error to the 16S copy number measurements from the mouse gut dataset, limiting our analyses to the stool samples where copy numbers varied by an order of magnitude. Across 50 iterations, each copy number could randomly vary by a given percentage of error in either direction. We re-calculated *β*-diversity (*BC*^*A*^ and *GU*^*A*^ at *α* = 1 and *α* = 0.2) and compared these measurements to the original dataset.

Introducing random variation into measured 16S copy number altered *GU*^*A*^ values only modestly, and the effect was proportional to the magnitude of the added noise (Fig. 4C–D). At α = 1, each 1% of quantification error introduced an average difference of 0.0022 in *GU*^*A*^; at α = 0.2, *GU*^*A*^ was even less sensitive. Thus, moderate α values provide a balance between interpretability and robustness to noise in absolute quantification. The max deviation from true that added error could inflict on a given metric was also proportional (and always less) than the magnitude of the error itself (Fig. 4D). A more rigorous approach to assessing error propagation within these metrics, including mathematical proofs of the relationships estimated above, is outside the scope of this paper but would be helpful.

### Ecological Interpretation and conceptual significance

Absolute UniFrac reframes the interpretation of *β*-diversity by making biomass an explicit ecological axis rather than an unmeasured or normalized-away quantity. Conventional UniFrac captures composition and shared evolutionary history, but implicitly invites interpretation as if it also encodes differences on microbial load. By incorporating absolute abundance directly, Absolute UniFrac helps to resolve this mismatch and restores alignment between ecological hypotheses and the quantities represented in the metric. In this view, β-diversity becomes a three-axis ecological measure, incorporating composition, phylogeny and biomass, rather than a two-axis approximation. That said, the additional dimension of microbial load also increases the complexity of applying and interpreting this metric.

There are many cases where the incorporation of absolute abundance allows microbial ecologists to assess more realistic, ecologically-relevant differences in microbial communities, especially when microbial load is mechanistically central. Outside of the datasets re-analyzed here [3, 5, 9, 10]; the temporal development of the infant gut microbiome involves both a rise in absolute abundance and compositional changes [6]; bacteriophage predation in wastewater bioreactors can be understood only when microbial load is considered [20]; and antibiotic-driven declines in specific swine gut taxa were missed using relative abundance approaches [21]. As *β*-diversity metrics (and UniFrac specifically) remain central to microbial ecology, these findings highlight how interpretation changes once biomass is incorporated. Broader adoption of absolute abundance profiling will also depend on data availability. Few studies currently make absolute quantification data publicly accessible, underscoring the need to deposit absolute measurements alongside sequencing reads for reproducibility, ideally as metadata within SRA submissions.

While demonstrated here with 16S rRNA data, the approach should extend to other marker genes or (meta)genomic features, provided absolute abundance estimates are available. In this sense, *GU*^*A*^ offers a more ecologically grounded view of lineage turnover by jointly reflecting variation in biomass and phylogenetic structure.

No single metric (or *α* in *GU*^*A*^), however, is universally “best”. Each β-diversity metric emphasizes a different dimension of community change. Researchers should therefore select metrics based on the ecological quantity that is hypothesized to matter most (Box 1). Here, we demonstrate not that *GU*^*A*^ outperforms other measures, but that it faithfully incorporates the three axes of variation it was designed to incorporate: composition, phylogenetic similarity, and absolute abundance (Fig. 1 and Fig. 2). As with any *β*-diversity analysis, interpretation requires matching the metric to the ecological question at hand, and exploring sensitivity across different metrics where appropriate [22]. By providing demonstrations and code for the application and interpretation of *U*^*A*^*/GU*^*A*^, we hope to encourage the use of these metrics as a tool of microbial ecology.

## Conclusion

By explicitly incorporating absolute abundance, Absolute UniFrac shifts phylogenetic β-diversity from a two-axis approximation to a three-axis ecological measure. This reframing connects the metric to the underlying biological questions that motivate many microbiome studies. As methods for quantifying microbial load continue to expand, the ability to interpret β-diversity in a biomass-aware framework will become increasingly important for distinguishing true ecological turnover from proportional change alone. In this way, Absolute UniFrac is not simply an alternative distance metric but a tool for aligning statistical representation with ecological mechanism.

**Box 1: β-diversity metrics should reflect the hypothesis being tested**

Absolute UniFrac is most informative when variation in microbial load is expected to carry ecological meaning rather than being a nuisance variable. The choice of α determines how strongly abundance differences influence the metric, and should therefore be selected based on the hypothesis, not by convention. In settings where biomass is central to the mechanism under study, higher α values appropriately foreground that signal, whereas in cases where load variation is incidental or confounding, lower α values maintain interpretability. Framing α as a hypothesis-driven choice repositions β-diversity from a default normalization step to an explicit ecological decision.

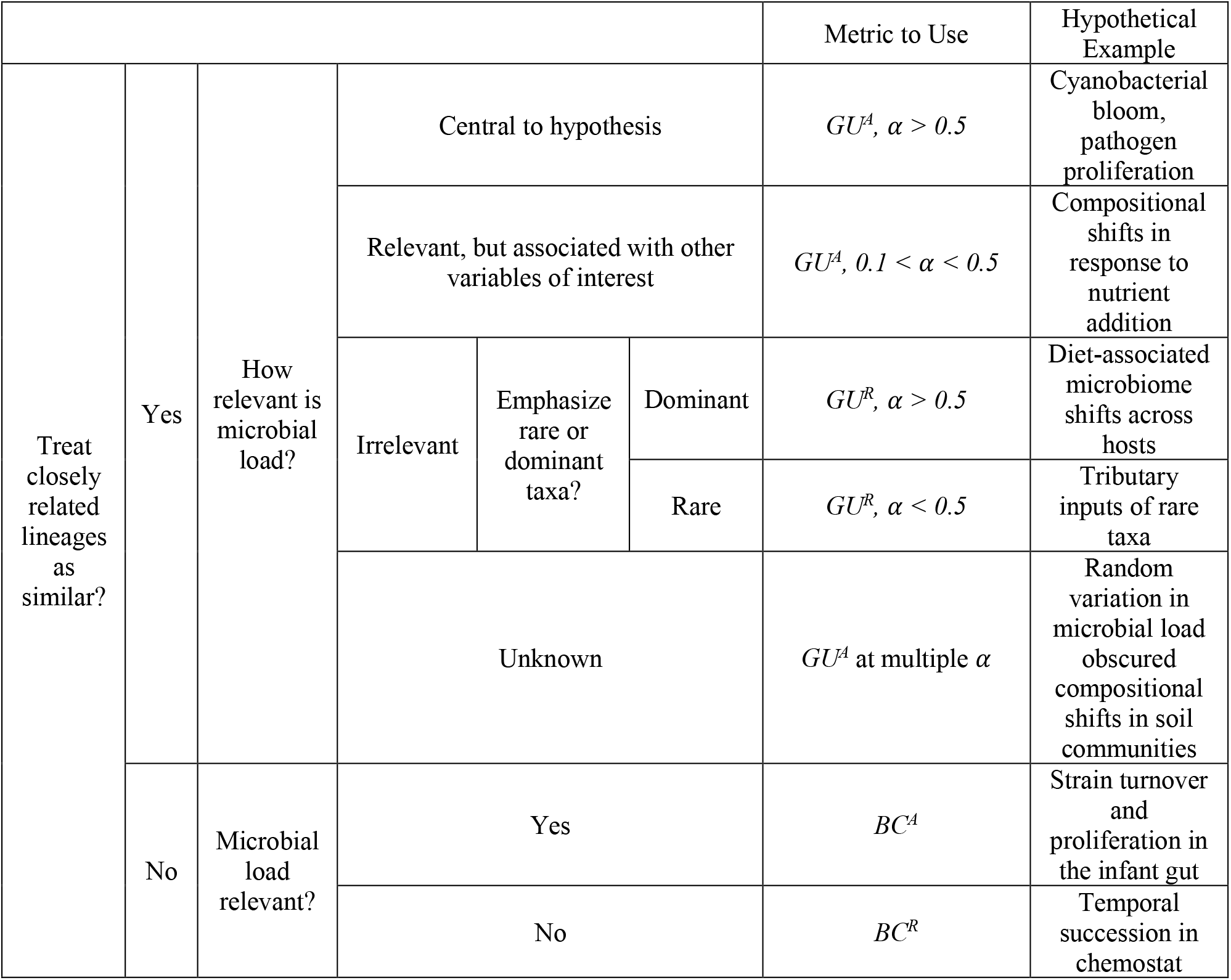

**Box 2: Rarefaction workflow for incorporating absolute abundance**

While we refrain from an in-depth analysis of rarefaction approaches, here we present our workflow for incorporating rarefaction alongside absolute abundance. First, samples were assessed for anomalously low read counts and discarded (sequencing blanks and controls were also removed). For rarefaction, each sample in the ASV table was subsampled to equal *sequencing* depth (# of reads) across 100 iterations, creating 100 rarefied ASV tables. These tables were then converted to relative abundance by dividing each ASV’s count by the equal sequencing depth (rounding was not performed). Then, each ASV’s absolute abundance within a given sample was calculated by multiplying its relative abundance by that sample’s total cell count or 16S copy number. Methods to predict genomic 16S copy number for a given ASV were not used [23].

Each distance/dissimilarity metric was then calculated across all 100 absolute-normalized ASV tables. The final distance/dissimilarity matrix was calculated by averaging all 100 iterations of each distance/dissimilarity calculation. Of note for future users: pruning the phylogenetic tree after rarefaction for each iteration is not necessary – ASVs removed from the dataset do not contribute nor change the calculated of UniFrac distances.

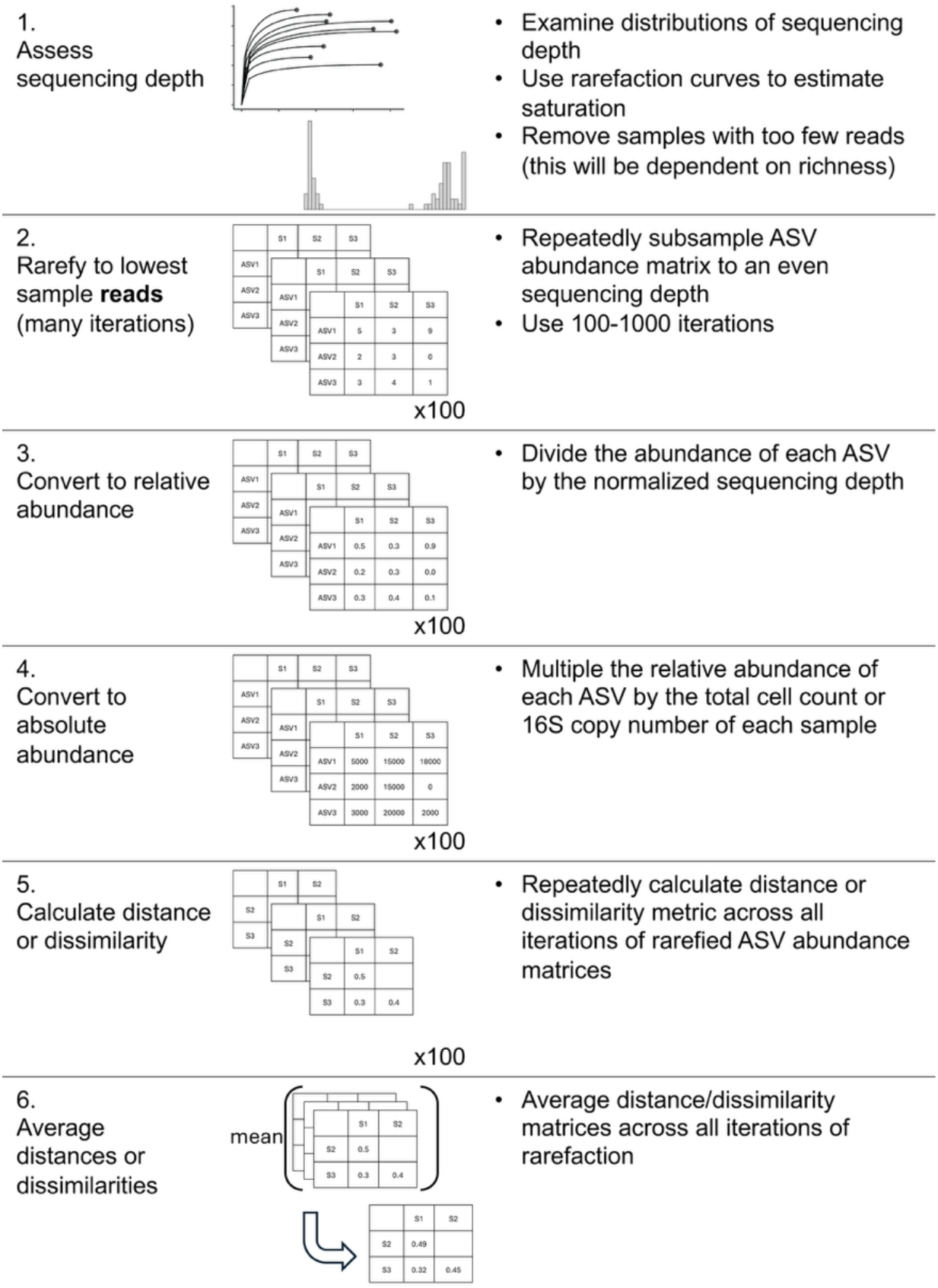

