## Supporting_Information for "Interpreting UniFrac with Absolute Abundance: A Conceptual and Practical Guide"

### Supporting Methods

*ASV Generation and Phylogenetic Tree Construction*

Sequencing data and identifying metadata were downloaded from the Sequence Read Archive (SRA), from BioProject IDs PRJNA815056, PRJNA575097, PRJNA1212049, and PRJNA302180 [1–4]. Full details and code of the Pendleton et al. 2025 data analysis are included within that paper and associated Github repository and not shown here. Each dataset varied substantially in terms of which 16S region it targeted, sequencing strategy, and read quality, so ASV generation varied between them in terms of primer removal, filtering, and trimming (see code for full description of these steps). Post trimming, all ASVs were generated using the same methods within the standard DADA2 workflow [5]. Chimaeras were removed, and ASVs were size selected (252/253 bp for V4 datasets, >400bp for V3-V4 datasets). Taxonomy was assigned via the Silva v138.2 database, and used to remove mitochondrial and chloroplast sequences [6]. When sequencing positives or negatives were present, they were removed.

Phylogenetic trees were built using alignment via MAFFT followed by FastTree under a generalized time-reversible model [7, 8]. Trees were visualized via ggtree in R, and anomalously long branches were removed using ape [9]. Trees, metadata, taxonomy, and ASV abundances (OTU tables) were organized and analyzed using phyloseq [10].

*Rarefaction and $\beta$-diversity*

To generate rarefied ASV tables of equal sequencing depth, ASV abundance matrices were subsampled using a multivariate hypergeometric distribution via the rmvhyper function in the extraDistr package (see generate_rarefied_abs_tables.R) [11]. Each ASV was then converted to relative abundances, and then to absolute abundances by multiplying the relative abundance by each samples cell count or 16S copy number. Bray-Curtis dissimilarities were calculated via the vegdist function in vegan [12]. Unless otherwise noted, all Unifrac distances were calculated via the GUnifrac package [13]. Final distance matrices were the average of all rarefied distance matrices. All samples within each dataset were used for contour plots in Figure 2.

*PERMANOVAs and Ordinations*

PERMANOVAs were conducted via the adonis2 function in vegan (Fig. 3 and Fig. S2). To limit confounding variables, not all samples were used in these analyses. From the cooling water dataset, just samples from Reactor cycle 1 were used. For the mouse gut, just stool samples were used. For the soil dataset, just mature samples were used. All PERMANOVAs were run with 1,000 iterations. These same, simplified datasets were used for Principal Coordinates Analysis in Fig S3.

*Timing Analysis*

To estimate computational time, we subsampled the soil dataset to a set number of ASVs, samples, and $\alpha$ numbers. When testing ASV number, we used 10 samples and one $\alpha$ value, when testing sample or $\alpha$ values, ASVs were held constant at 2,000. Each case was replicated 20 times, and computation time was calculated via the microbenchmark function from the microbenchmark package, with two replications each time [14].

*Error Analysis*

To estimate the impact of random error on quantification methods, we used the mouse gut dataset, focusing only on the stool samples. These samples ranged in 16S copy number from $10^{11}$-$10^{12}$ copies/gram. We tested a range of potential error, from 1% up to 50%. For each error percentage, the amount of error was selected from a normal distribution with a mean of that error percent and a standard deviation 1/10$^{th}$ of that error percentage. This error was then randomly assigned a direction (by multiplying by a binomial distribution of -1 and 1), and multiplied by the copy number to create a deviation from the true value, which was added to the original value. For example, in the case of 50% error, we first drew a random selection of error values from a distribution with mean 0.5 and standard deviation of 0.05. These errors were then randomly assigned to be negative or positive, and multiplied by the original cell counts, plus the cell count itself. We repeated this fifty times. Across these fifty iterations, we first rarefied the ASV abundances to relative abundance and then normalized to absolute abundance using these error-added values. We then compared the absolute difference in $GU^A$ or $BC^A$ from these error-added datasets compared to the original data to produce Fig. 4C-D.

*Other Coding Packages*

Other packages used for general coding and visualization include tidyverse, purr, patchwork, NatParksPalette, broom, corrr, ggpubr, and renv. All packages and version numbers are listed in Table S1.

### Supplemental Figures

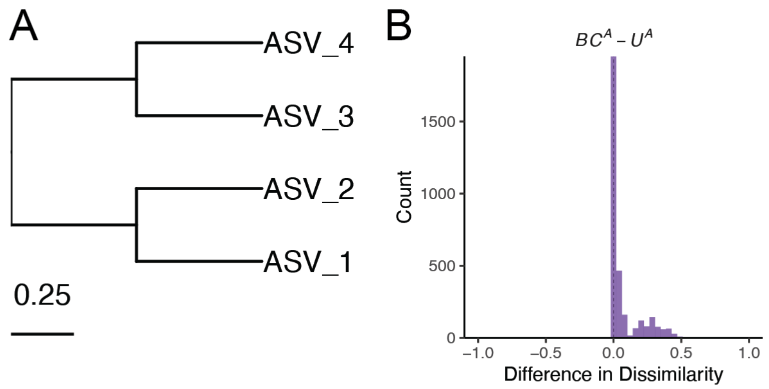


*Figure S1.* $U^A$ is always less than $BC^A$ when branch lengths are fully symmetrical. (A) Symmetrical tree used for simulations as opposed to non-symmetrical tree in Fig. 1A. (B) Distribution of differences between $BC^A$ and $U^A$. As the differences are never negative, $U^A$ is always less than or equal to $BC^A$.

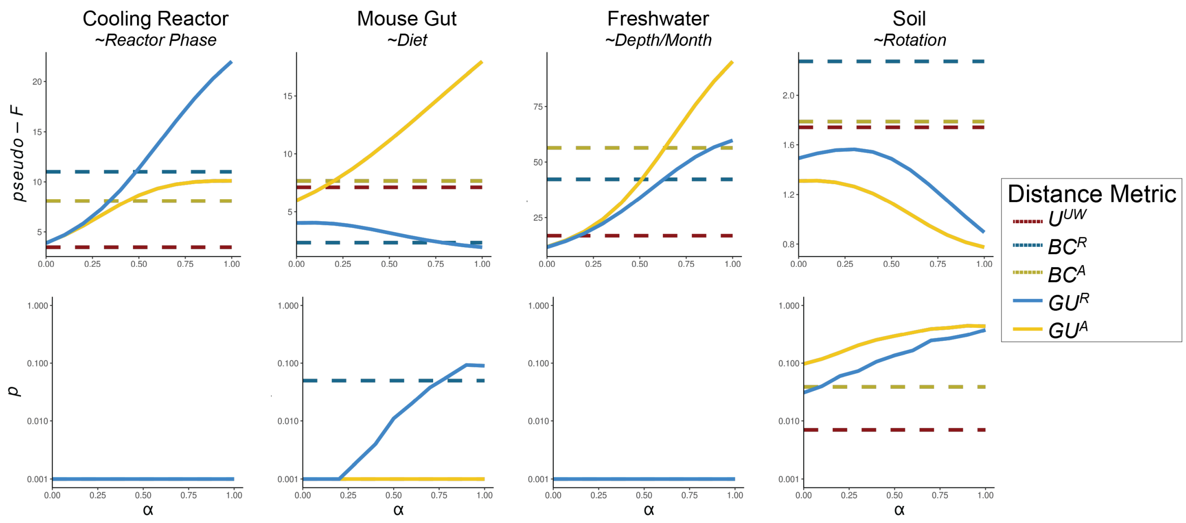


*Figure S2. Additional PERMANOVA results when using $GU^A$ across a range of α values.* PERMANOVAs were run testing the significance of two-three category groups from each dataset (provided in italics beneath data names). Results indicate *pseudo-F* statistics and *p*-values after 1,000 iterations. In the cooling reactor, only samples from Reactor cycle 1 were used; in the mouse gut, only stool samples were used, and in the soil, only mature samples were used. Note the y-axes for *pseudo-F* plots are variable between datasets, and y-axes for the *p*-value plots are log-scaled.

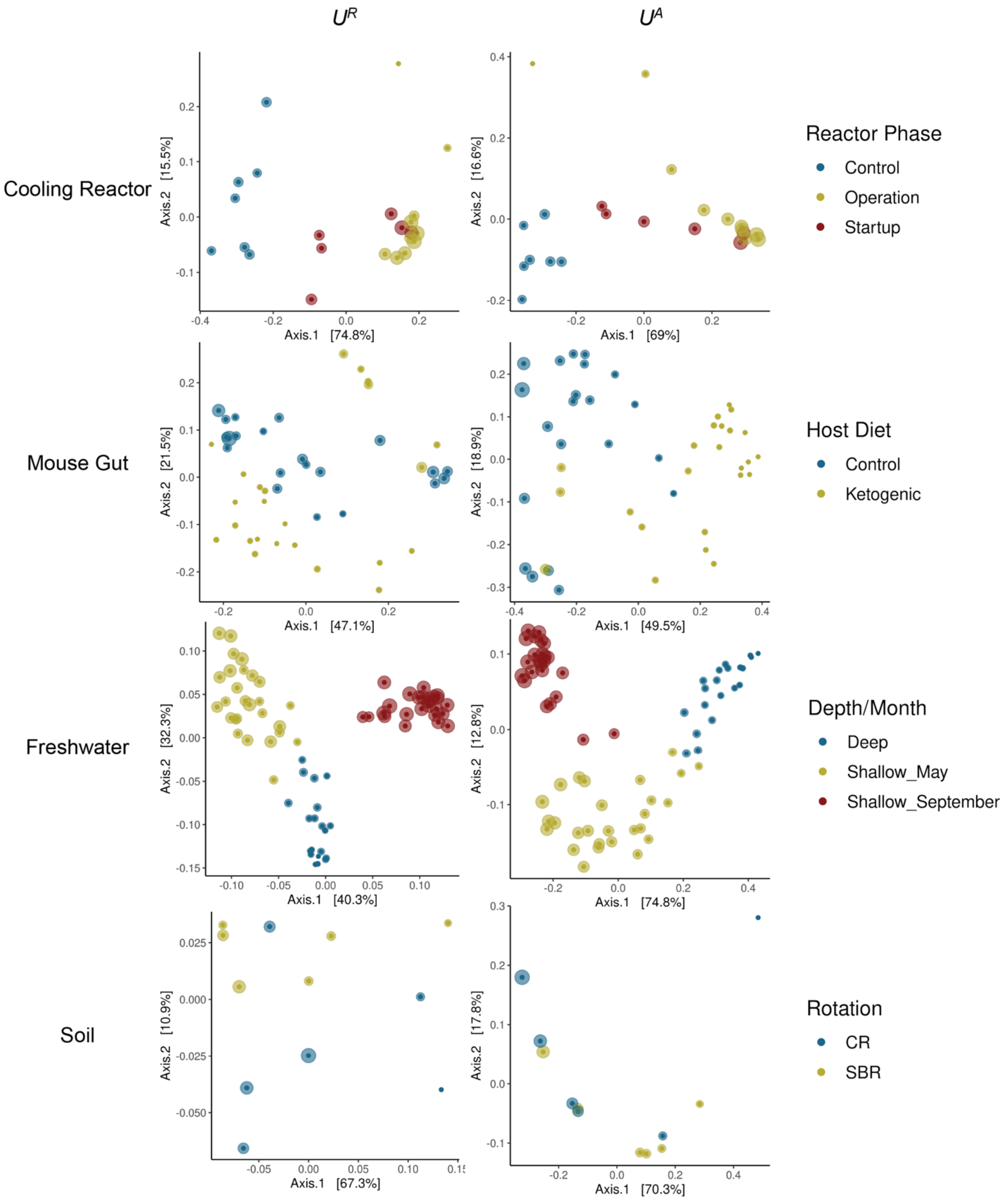


*Figure S3. Principal Coordinate Analysis ordinations of each dataset using $U^R$ and $U^A$.* Points are colored using the same categorical variable tested in the PERMANOVAs of Fig. 2 and Fig. S2 (for additional details on experimental design, see [1, 3, 4, 15]). Both $U^R$ and $U^A$ were calculated at an $\alpha = 1$.

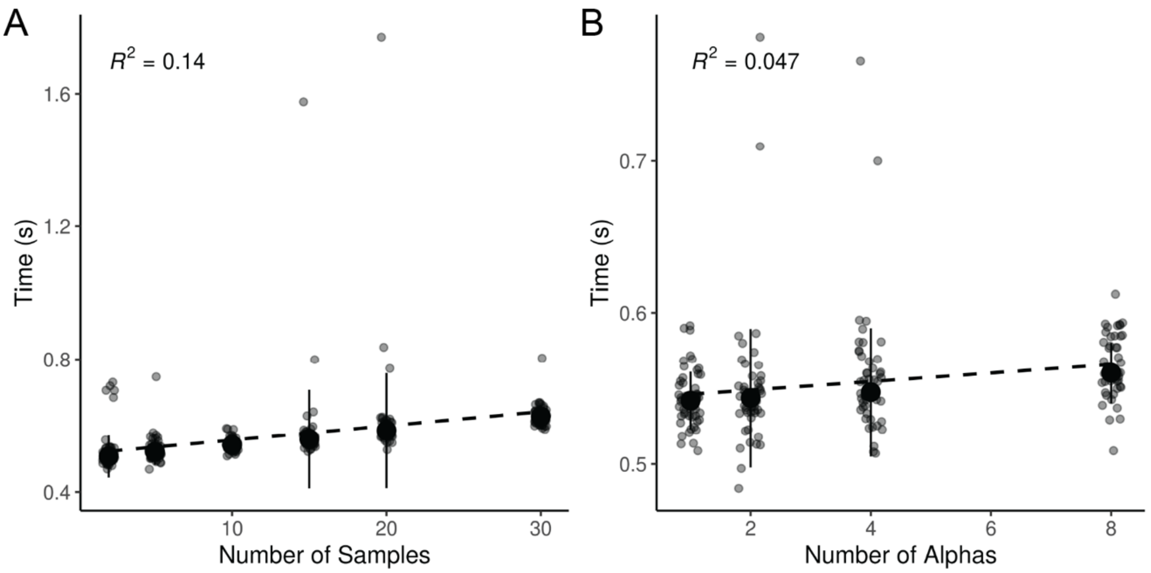


*Figure S4. Additional parameters which weakly influence computation time for $GU^A$.* A) $GU^A$ was calculated 50 times across six sample sizes (2, 5, 15, 20, 30) with a constant of 2,000 ASVs and one calculated $\alpha$ (though unweighted Unifrac is also calculated by default). B) $GU^A$ was calculated 50 times across four alpha parameter sizes (1, 2, 4, 8; note unweighted Unifrac is also calculated by defualt) with a constant of 2,000 ASVs and 10 samples. In both panels, $R^2$ is derived from a linear model between the x and y axes.

| Package/Software | Version | Citation |
| --- | --- | --- |
| R | 4.5.0 | [16] |
| RStudio | 2024.12.1+563 | [17] |
| tidyverse | 2.0.0 | [18] |
| phyloseq | 1.52.0 | [10] |
| vegan | 2.7-1 | [12] |
| GUniFrac* | 1.8.1 | [13] |
| ggtree | 3.16.0 | [19] |
| patchwork | 1.3.1 | [20] |
| NatParksPalettes | 0.2.0 | [21] |
| ape | 5.8-1 | [9] |
| broom | 1.0.8 | [22] |
| corrr | 0.4.4 | [23] |
| renv | 1.0.5 | [24] |
| microbenchmark | 1.5.0 | [14] |
| ggpubr | 0.6.1 | [25] |
| dada2 | 1.36.0 | [5] |
| MAFFT | 7.520 | [7] |
| FastTree | 2.1.11 | [8] |
| cutadapt | 5.1 | [26] |
| extraDistr | 1.10.0 | [11] |

*Table S1. Software and packages used in analysis.* Note that GUniFrac was modified slightly to make incorporating absolute abundances more apparent; this version can be installed via Github at https://github.com/MarschmiLab/GUniFrac.
